# Evolution of the acoustic startle response of Mexican cavefish

**DOI:** 10.1101/809665

**Authors:** Alexandra Paz, Brittnee McDole, Johanna E. Kowalko, Erik R. Duboue, Alex C. Keene

## Abstract

The ability to detect threatening sensory stimuli and initiate an escape response is essential for survival and under stringent evolutionary pressure. In diverse fish species, acoustic stimuli activate Mauthner neurons, which initiate a stereotypical C-start escape response. This reflexive behavior is highly conserved across aquatic species and provides a model for investigating the neural mechanism underlying the evolution of escape behavior. Here, we define evolved differences in the C-start response between populations of the Mexican cavefish, *Astyanax mexicanus*. Cave populations of *A. mexicanus* inhabit in an environment devoid of light and macroscopic predation, resulting in evolved differences in diverse morphological and behavioral traits. We find that the C-start is present in multiple populations of cavefish and river-dwelling surface fish, but response kinematics and probability differ between populations. The Pachón population of cavefish have an increased response probability, a slower response and reduction of the maximum bend angle, revealing evolved differences between surface and cave populations. In two other independently evolved populations of cavefish, the response probability and the kinematics of the response differ from one another, as well as from surface fish, suggesting the independent evolution of differences in the C-start response. Investigation of surface-cave hybrids reveals a relationship between angular speed and peak angle, suggesting these two kinematic characteristics are related at the genetic or functional levels. Together, these findings provide support for the use of *A. mexicanus* as a model to investigate the evolution of escape behavior.

## Introduction

Predator evasion is essential for survival and is thought to be a critical trait contributing to behavioral adaptation in novel environments (Domenici, 2010). Multiple sensory systems are used to detect predators including olfaction, vision, and mechanotransduction, which all result in the activation of arousal systems (Ferrari et al, 2010; Bleicher et al, 2018; Temizer et al, 2015; Franceschi et al, 2016; Mooney et al, 2016; Suzuki, 2018). The escape responses of a variety of larval fish systems have been studied in detail, including zebrafish, medaka, killifish, and goldfish (Burgess & Granato, 2007; Featherstone, 1991; Canfield, 2006; Fleuren et al, 2018). All of these species exhibit a conserved, highly stereotypical C-start response. The startle response of fish is termed the C-start because of the characteristic c-shaped curve formed by the body during the first stage of the response, which is followed by a smaller counter-bend and rapid swimming (Kalueff et al, 2013). It is also highly stereotyped and plastic, providing a system to examine innate behaviors and their experience-dependent modification (Lopez-Scheir, 2016).

The escape responses of larval fish are initiated by multiple pairs of highly conserved reticulospinal neurons that receive input from a variety of sensory systems and project to spinal interneurons and motor neurons that innervate the muscles of the trunk (Liu & Fetcho, 1999; Gahtan et al, 2002; Kohashi & Oda, 2008; Bosch & Paul, 1993). Activation of one of these pairs of neurons, the Mauthner cells, initiate a stereotype short latency C-start escape reflex (Burgess & Granato, 2007; Liu & Fetcho, 1999). Mauthner cells receive input from multiple sensory modalities including from the visual, olfactory, and mechanosensory systems (Medan et al, 2018; Kohashi & Oda, 2008; Canfield, 2006; Kimmel et al, 1990; Bhattacharyya et al, 2017). Thus, these neurons receive sensory information and initiate escape reflexes, providing a model for investigating sensory-motor integration (Bierman et al, 2009). Despite its fundamental importance to behavioral evolution, surprisingly little is known about the neural mechanisms through which ecological perturbation shapes the evolution of this escape response.

The Mexican cavefish, *Astyanax mexicanus* is a powerful model for studying behavioral evolution (Keene, McGaugh, & Yoshizawa, 2015; Gross, 2012). These fish exist as surface fish that inhabit rivers in Mexico and Southern Texas and at least 29 geographically isolated cave-dwelling populations of the same species (Mitchell, Russell, & Elliott, 1977; Jeffery, 2009). The ecology of caves differs dramatically from the surface habitat resulting in the emergence of distinct morphological and behavioral phenotypes. For example, the absence of light in caves is thought to contribute to the evolution of albinism, eye-loss, and circadian rhythm (Keene et al, 2015). As a consequence of these environmentally driven changes, these fish are useful models for investigating convergent trait evolution, and more recently, the evolution of neural circuits mediating behavior (Jaggard et al, 2018; Alie, 2018; Duboué, 2012). Interestingly, no macroscopic predator the caves lack macroscopic predators of *A. mexicanus*, raising the possibility that a lack of selective pressure for predator avoidance contributes to morphological and behavioral evolution in cavefish populations (Pitcher, 1986).

Prominent changes in sensory processing contribute to behavioral evolution in cavefish. This includes enhanced sensitivity of the lateral line that contributes to prey capture and sleep loss in cavefish (Yoshizawa et al, 2012; Lloyd et al, 2018; Jaggard et al, 2017). Cavefish have also evolved increased sensitivity to tastants and odorants, presumably to support efficient foraging in the absence of visual cues (Shiriagin & Korsching, 2019; Bibliowicz, 2013; Hinaux et al, 2016). Additionally, *A. mexicanus* use acoustic stimulation to communicate, and a recent report highlights the differences in this communication between surface and cave morphs (Hyacinthe et al, 2019). The diversity of evolved changes in sensory processing combined with the robust ecological differences raises the possibility that the startle reflex may differ between populations of *A. mexicanus*.

Here, we systematically investigate the evolution of the C-start response to acoustic stimuli in multiple *A. mexicanus* population. We find differences in both response probability and kinematics between surface fish larvae and three different populations of cavefish. These findings support the notion that the ecological differences between cave and river environments contribute to differences in escape behavior and provide a platform for investigating the evolution of neural circuits contributing to sensory-motor integration.

## Results

To quantify differences in startle response, we constructed a system to produce acoustic pulses similar to those shown to induce startle behavior in zebrafish (Burgess & Granato, 2007; Bhandiwad, 2013; Zeddies & Fay, 2005). Fish were individually placed in custom-designed wells attached to a vibration exciter that provided acoustic stimuli. Behavior of the fish was recorded throughout the stimulation using a high-speed camera (Fig 1A). We first compared the probability of 6 day post fertilization (dpf) surface fish and Pachón cavefish initiating a C-start in response to acoustic stimulation. We found that both surface and Pachón cave populations responded to acoustic stimuli with a stereotyped response consisting of simultaneous head and tail turning, as observed during classic C-start escape reflexes that have been characterized in zebrafish and other aquatic models (Burgess & Granato, 2007; Featherstone, 1991; Canfield, 2006; Fleuren et al, 2018). To determine whether the sensitivity required to elicit an escape response differed between populations, we quantified the probability of C-start initiation in surface fish and Pachón cavefish at multiple vibration intensities and found that cavefish exhibit an increased response probability to vibrations at 31 dB (surface fish 67%, cavefish: 53%) and 35 dB (surface fish 74%, cavefish: 90%), but not 28 dB (surface fish: 47%, cavefish: 43%) (Fig 1B). These data demonstrate that Pachón cavefish have a more acute sensitivity to vibrations relative to their surface conspecifics.

**Figure 1.**
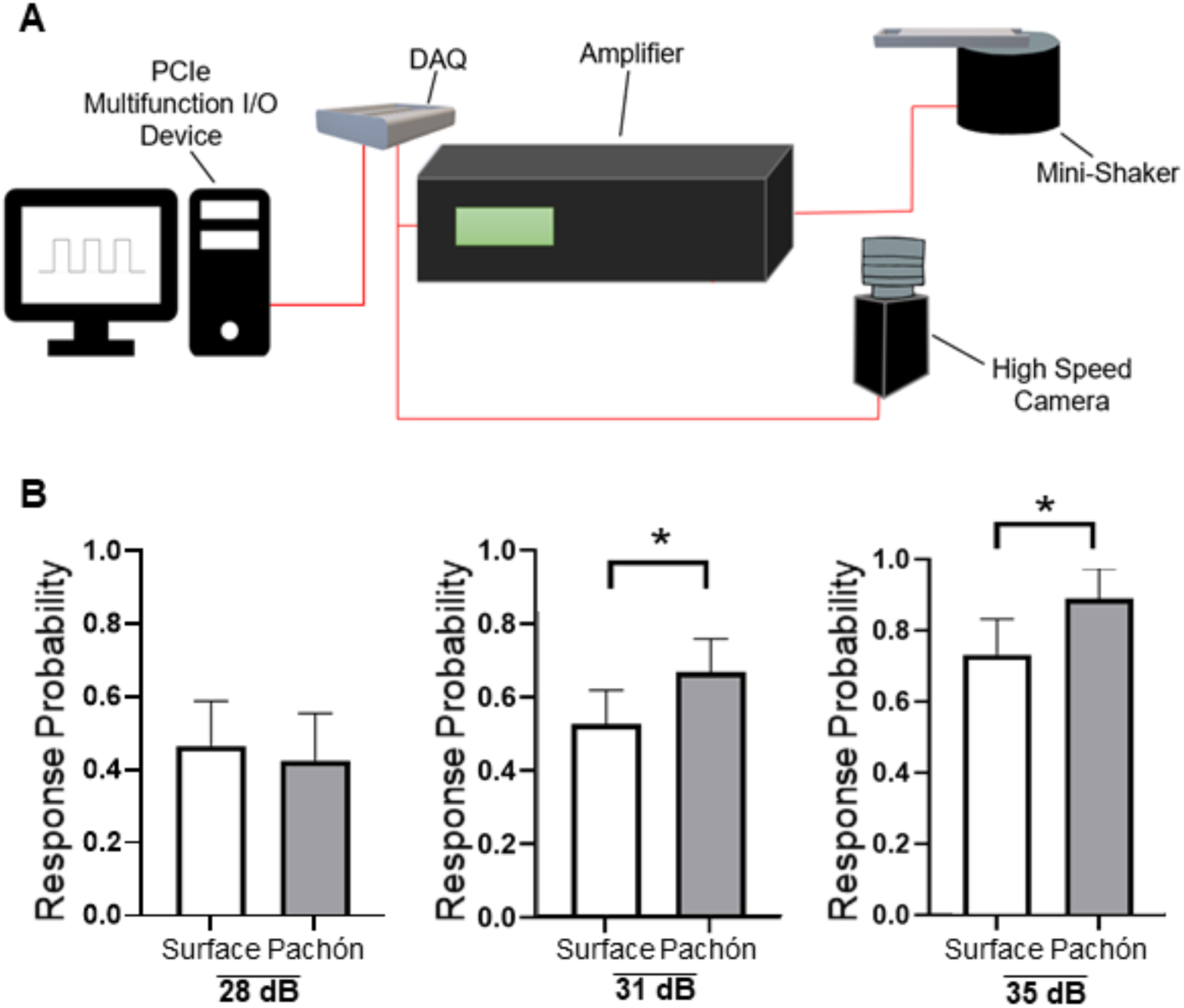
Measurements of C-start response in *A. mexicanus*. (A) Acoustic stimuli were generated using an amplifier and small vibration exciter controlled by a Data acquisition device (DAQ). A high-speed camera collected data throughout the stimulation protocol. (B) Pachón larvae (grey) exhibit increased startle probability to vibrational stimuli at intensities of 31 dB (SF N=112, Pa N=103, 2-tailed Fisher’s Exact Test p=0.0374) and 35 dB (SF N=72, Pa N=48, 2-tailed Fisher’s Exact Test p=0.0373) compared to surface fish (white). No significant differences were detected at 28 dB (SF N= 64, Pa N= 56, 2-tailed Fisher’s Exact Test p=0.715). Error bars signify margin of error. * denotes p≤0.05.

In order to compare C-start kinematics between surface fish and cavefish, we quantified response latency, maximum change in orientation (referred to as “peak bend angle”), and angular speed, and found that the responses of surface fish and Pachón cavefish differ in all quantified kinematic parameters (Fig 2A & B). The C-start responses of Pachón cavefish are characterized by a decrease in angular speed and peak bend angle compared to surface fish, with Pachón turning approximately 3°/ms more slowly and to a peak bend angle that is smaller in magnitude by almost 20° relative to surface fish (Fig 2C & D). Pachón larvae also displayed significantly longer response latencies than surface fish larvae (Fig 2E). In surface fish, the shortest latency C-starts were initiated 7-9 ms after stimulus onset, in contrast to Pachón larvae in which the shortest latency C-starts were initiated 11-13 ms after stimulus onset (Fig 2F). Together these data suggest that cavefish have developed substantial differences in the C-start response.

**Figure 2.**
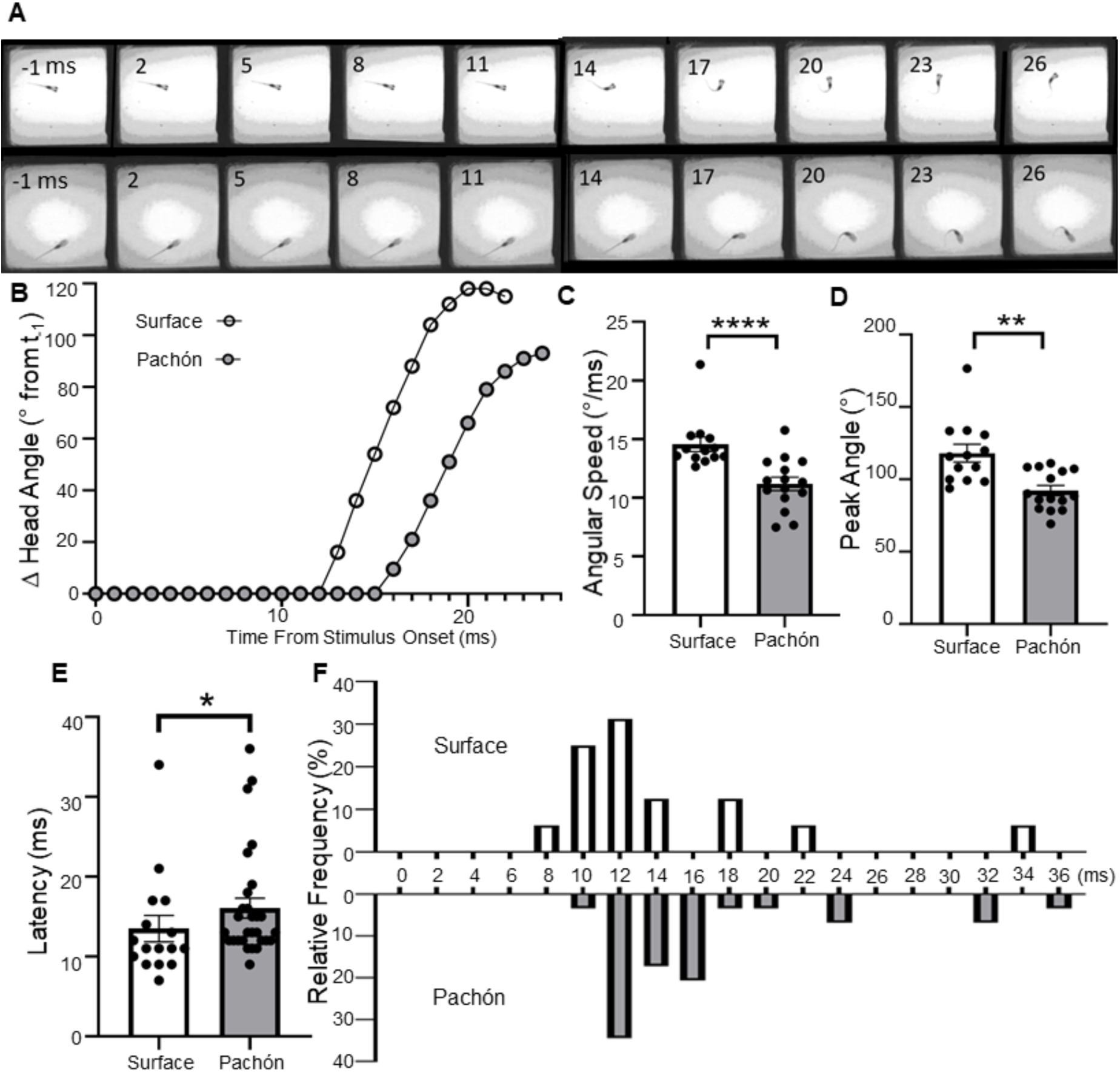
C-start kinematics are altered in Pachón cavefish. (A) Time lapse images showing typical surface fish (top) and Pachón cave fish (bottom) C-start responses. Changes in orientation over the course of the response were standardized to the fish’s orientation 1 ms before stimulus onset (first frame shown). Snapshots shown are 3 ms apart (B) Surface fish (open circles) and Pachón cavefish (filled circles) exhibit robust differences in c-start kinematics. Quantitative analysis was done to compare the angular speed, peak bend angle, and latency of surface and cave fish responses. (C) Comparisons of Pachón (N=15, median=11.34°/s) and surface fish (N=13, median=14.01°/s) responses revealed that cavefish exhibit significantly reduced turning speed than surface fish. Mann-Whitney U=17, p<0.0001. (D) Pachón cavefish (N=15, median=88.36°) also exhibit a smaller change in orientation during the first phase of the C-start response than surface fish (N=13, median 115.8°). Mann-Whitney U=29.50, p=0.0011. (E) Initiation of Pachón responses (N=29, median=13ms) was delayed relative to surface fish responses (N=16, median=11ms). Mann-Whitney U=145.5, p=0.0386. (F) A histogram of response relative frequency across different response latencies reveals a shift in Pachon caveifhs (black) to slower response latency. Error bars denote std. error of mean. * denotes P≤0.05. ** denotes P≤0.01, **** denotes P≤0.0001.

In teleosts, Mauthner neurons integrate visual stimuli, and a loom stimulus is enough to initiate a C-start response (Temizer et al, 2015; Bhattacharyya et al, 2017). Cavefish can detect light and sense looming stimuli, despite eye degeneration, raising the possibility that light modulates the C-start response (Yoshizawa & Jeffery, 2008). To assess the influence of visual input on response probability and kinematics we assayed escape response under light and dark conditions. The presence of light had no effect on response probability, response latency, or angular speed in cavefish or surface fish (Fig 3A-C). In goldfish, it was found that peak C-start bend angle was predictable based off of a fish’s orientation relative to the startle-inducing stimulus, except for situations where the predicted trajectory was blocked by a wall (Eaton & Emberley, 1991). This trend was true even when C-starts were initiated from rest, precluding the possibility that the lateral line was influencing escape kinematics. Furthermore, in zebrafish it has been shown that peak bend angle is a reliable predicter of escape trajectory (Bhattacharyya et al, 2017). In dark conditions, surface fish display an increase in peak bend angle, while no difference is detected in cavefish (Fig 3D). These data suggest that, as with goldfish, the escape path of surface fish is visually informed.

**Figure 3.**
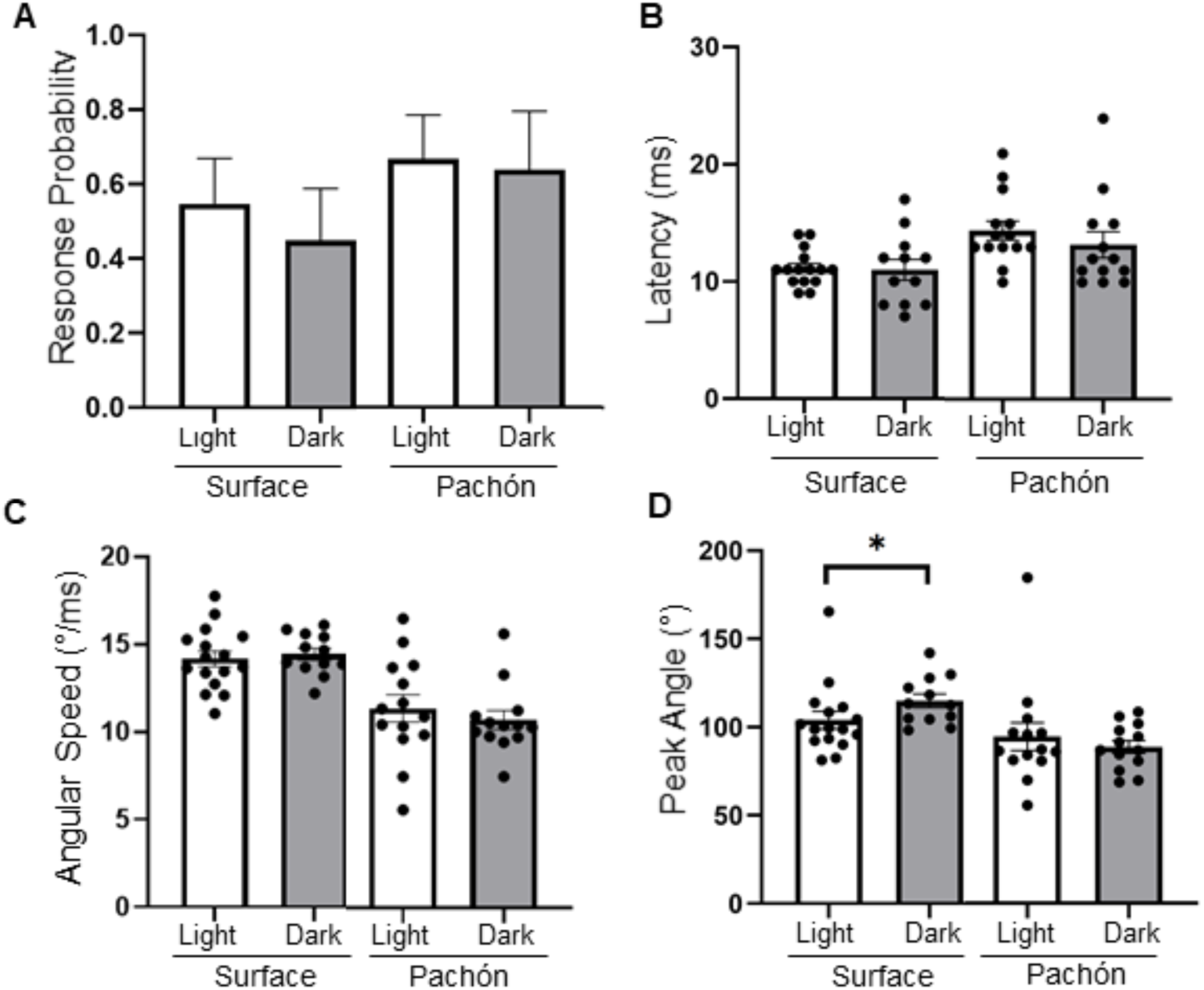
Visual input influences escape trajectory of C-start responses in surface fish. (A) The response probabilities of both surface fish (unfilled bars; light N=64, dark N=49) and Pachón cavefish (filled bars; light N=60, dark=36) were unaffected by the presence or absence of white light. Error bars signify margin of error. Surface fish: Fisher’s Exact test p = 0.345; Pachón: Fisher’s Exact test p=0.827. (B) The Latency of surface fish (light N=15, dark N=12) and Pachón cavefish (light N=14, dark N=13) responses also were unaffected. Surface fish unpaired t-test t = 0.1467, df = 25, p = 0.8846; Pachón unpaired t-test t = 0.8779, df = 25,p = 0.8779. (C) Likewise, the angular speed of surface fish (light N=16, dark N=12) and Pachón cavefish (light N=14, dark N=13) were unaffected by light. (D) The peak bend angle of surface fish (light N=16, dark N=12) was significantly larger in the absences of light. Median angle in light conditions was 99.15° and 112.8° in the dark. 2-tailed Mann-Whitney test U = 46.50, p = 0.0204. Pachón cavefish exhibited no difference in peak bend angle in dark conditions. Pachón 2-tailed Mann-Whitney test U = 87.50, p = 0.8774. * denotes p≤0.05. Error bars on kinematic data (B-D) signify standard error of the mean.

Independently evolved populations of cavefish have converged on numerous behavioral and morphological traits (Keene, McGaugh, & Yoshizawa, 2015; Gross, 2012), providing a powerful system for examining whether convergent traits arise through similar or distinct genetic mechanisms. To determine whether the changes in C-start probability and kinematics are shared across cavefish populations, we measured response in Molino and Tinaja cavefish. While Tinaja larvae exhibit a response probability similar to that of surface fish, larvae from the Molino population exhibited a 98% response probability, which was significantly higher than surface fish and any of the cavefish populations (Fig 4A). Unlike Pachón larvae, Molino and Tinaja did not exhibit any differences in response latency relative to surface fish (Fig 4B). However, angular speed was reduced in both Tinaja and Molino populations (Fig 4C) while the peak bend angle was significantly reduced in Tinaja compared to surface fish (Fig 4D). Together, these findings reveal convergence on a decrease in angular speed during the C-start response in cavefish. On the other hand, the variety in latency, peak bend angle, and response probability observed in the three cavefish populations analyzed here reveal the evolution of unique kinematic changes across all three cavefish populations.

**Figure 4.**
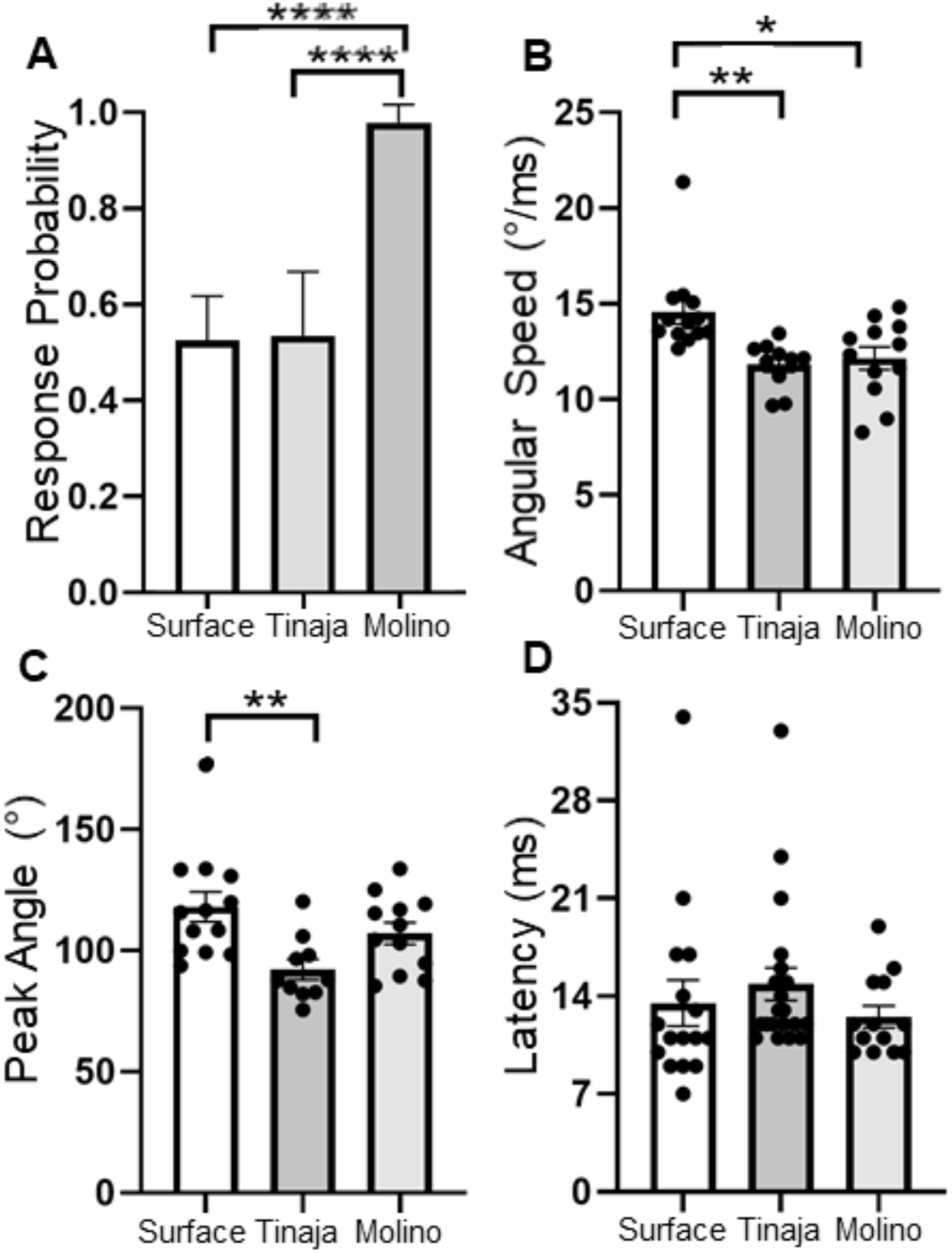
The C-start response is altered in Tinaja and Molino populations of cavefish. (A) Molino larvae (N=54) responded in 98% of trials, exhibiting significantly higher response probability than either surface (N=112) or Tinaja (N=54) larvae. Error bars denote margin of error. Surface/Molino X^2^ (1)=32.28, p<0.0001; Tinaja/Molino X^2^ (1)=26.292, p<0.0001. (B) There were no significant differences in response latency (surface N=16, Tinaja N=21, Molino N=13). One-way ANOVA F(2, 47) = 0.8153, p=0.4487. (C) Surface fish larvae (N=13) turned with significantly quicker angular speed than Tinaja (N=10) or Molino (12) larvae. One-way ANOVA F(2, 32)=0.7188, p=0.0024. Surface/Molino p=0.0101; surface/Tinaja p=0.0051. (D) Surface fish exhibited the most drastic change in orientation. One-way ANOVA F(2, 32)=0.7560. Surface/Tinaja p=0.0044. Error bars on kinematic data (B-D) denote standard error of the mean. * denotes p≤0.05, ** denotes p≤0.01, **** denotes p≤0.0001.

It is possible that independent genetic mechanisms contribute to different kinematic components of the C-start, or that they have evolved through shared genetic architecture. A benefit of *A. mexicanus* is that cavefish and surface fish populations are interfertile, producing hybrid offspring that possess behavioral and morphological characteristics ranging from cave-like to surface-like, as well as intermediate phenotypes. To differentiate between these possibilities, we quantified the kinematics of C-start responses of surface-cave F_2_ hybrid fish. The response probability of F_2_ hybrid fish was intermediate to pure surface and Pachón fish, but this did not reach significance (Fig 5A). Significant differences in latency were not detected between hybrid fish and surface or Pachón populations, with the range of values exhibited by F_2_ hybrids encompassing the full range of values seen in surface and Pachón fish. Though no significant differences were identified in these data, it is worth noting that the mean value for the F_2_ hybrids (16 ms) matched that of Pachón cavefish (16 ms), but not that of surface fish (14 ms) (Fig 5B). Similarly, the peak bend angle of F_2_ hybrids resembled those of Pachón cave fish, differing significantly from the larger bend seen in surface fish responses (Fig 5C). The angular speed of F2 hybrids was intermediate to that of surface and Pachón fish (Fig 5D). To determine the relationship between components of the C-start response, we quantified the correlation between each pair of kinematic parameters. No correlation was observed between response latency and peak angle or angular speed, however there was a significant correlation between angular velocity and peak angle. Taken together, these findings suggest that there are a variety of factors influencing the various kinematic parameters that compose the C-start response.

**Figure 5.**
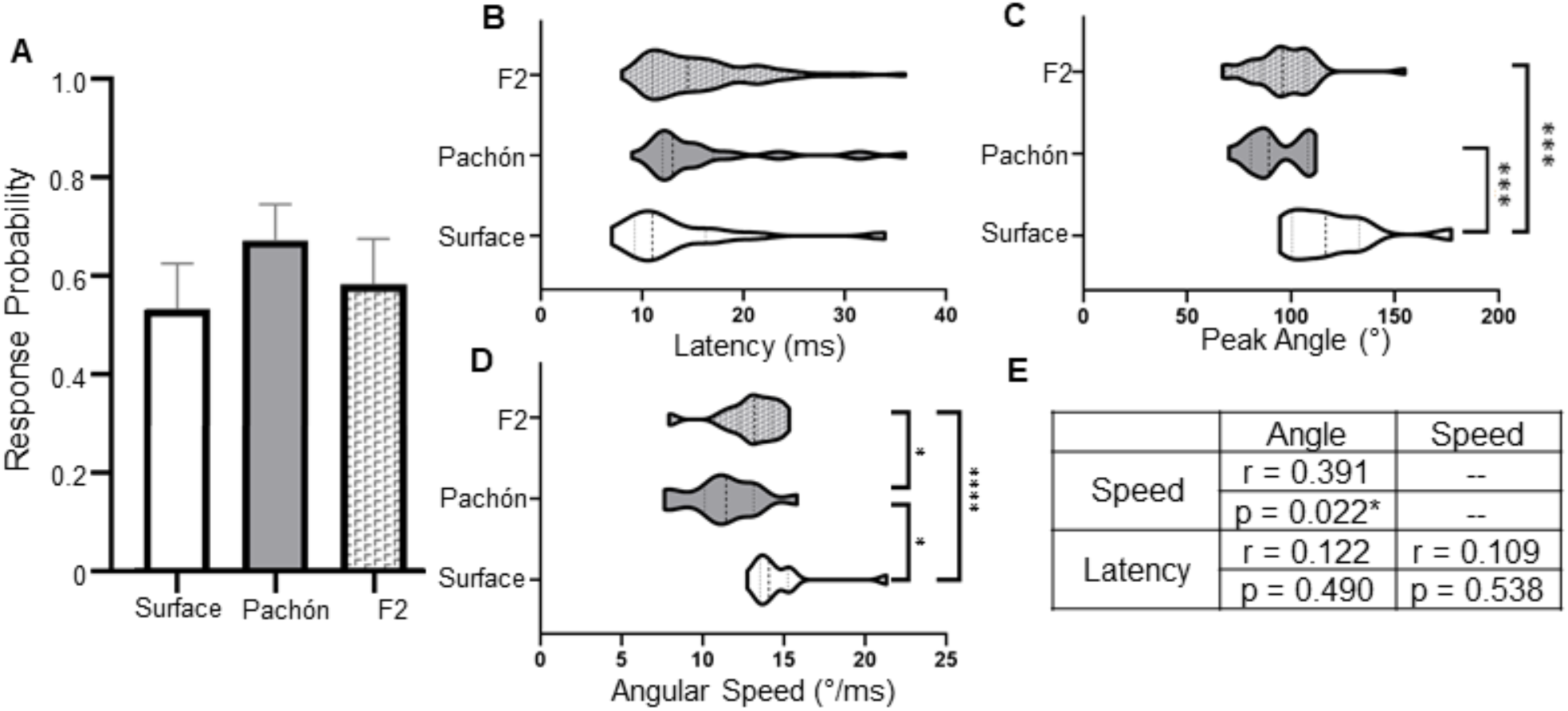
Analysis of surface-cave hybrids reveals a relationship between angular speed and peak C-start angle. (A) No difference was identified between the response probabilities of surface fish (N=112), Pachón (N=103), and F_2_ hybrids (N=179), X^2^ (2, N=291)=2.93, p=0.099, though surface and Pachón cavefish approached significance. Error bars signify margin of error (B) There was also no difference in response latency between surface (N=16), Pachón (N=29), and F_2_ hybrids (N=68). (C) F_2_ hybrids (N=34) exhibit a peak change in orientation similar to that of Pachón (N=15) larvae, in contrast to surface fish (N=13). One-way ANOVA F(2, 59) = 0.66, p < 0.001. Surface/Pachón p<0.001, surface/ F_2_ p<0.001. (D) The angular speed of the F_2_ hybrids (N=34) was intermediate to that of cavefish (N=15) and surface fish (N=13). One-way ANOVA F(2, 59) = 0.48, p = 0.0002. Surface/ F_2_ p=0.0426, F_2_/Pachón p=0.0139, surface/Pachón p<0.0001. (E) Spearman’s rank correlation test indicates a positive correlation exists between peak angle and speed. r denotes Spearman’s correlation coefficient, rho. Dotted lines in violin plots denote quartiles and median. * denotes p≤0.05, ** denotes p≤0.01, *** p≤0.001.

## Discussion

The C-start response represents a primary mechanism for predator avoidance in fish and amphibians (Yasugi & Hori, 2012; Walker et al, 2005; Fuiman, 1993) Here, we identify evolved changes in the C-start response in multiple independent populations of *A. mexicanus*. There are many differences between the ecology of caves and that of rivers and lakes inhabited by surface fish including changes in food availability, changes in water quality, loss of circadian cues, and reduced predation (Keene et al, 2015). It is possible that, since there is a near absence of predators in the caves, the changes observed in cavefish are due to relaxed interspecies selective pressure in the cave environments. However, adult surface and cave populations of *A. mexicanus* consume their larvae, raising the possibility that the C-start remains critical for intra-species predation. Further investigation of the ecology of early life environment within the natural setting may inform the cause of the evolved changes in the C-start.

In fish, escape responses can be characterized into those which occur quickly (short latency C-starts) and those that emerge later (long latency C-starts) (Burgess & Granato, 2007). Ablation of the Mauthner neurons completely abolishes short latency C-starts in goldfish and zebrafish, but not longer latency C-starts, which are initiated by a different set of reticulospinal neurons (Kohashi & Oda, 2008; Burgess & Granato, 2007; Eaton et al, 1982; Liu & Fetcho, 1999). We observed that the latency is significantly greater in Pachón cavefish than in surface fish, raising the possibility that differences in Mauthner neuron signaling may contribute to the observed differences in startle kinematics. In zebrafish, the frequency distribution of C-start initiation latency values produces a bimodal curve, with separate peaks representing short latency C-starts and less frequent long latency C-starts (Burgess & Granato, 2007; Takahashi et al, 2017; Issa et al, 2011). In our frequency analysis we did not identify separate peaks, however this is likely due to sample size, not a lack of long latency responses. Future studies testing a greater number of individuals may provide insight into how the reticulospinal escape network has evolved and the intrapopulation variation in this response.

We identified an increased probability of eliciting a startle response in Pachón and Molino larvae. It is possible that this is due to altered sensory detection of the acoustic stimuli, or due to changes at the level of processing that affect the threshold of Mauthner neuron activation. Adult surface and cavefish respond to click-like sounds that signal aggression, revealing the presence of acoustic communication between conspecifics in this species (Hyacinthe et al, 2019). Previous analysis of auditory sensitivity in *Astyanax* did not identify differences between surface and cave populations, supporting the notion that the differences observed are at the level of sensory processing, rather than detection (Popper, 1970, Hinaux, 2016). Therefore, it is unlikely that differences in sensory detection underlie the enhanced response probability in cavefish.

The kinematics of the C-start response differed between all three cavefish populations and surface fish. In Pachón and Tinaja cavefish, this is marked by a reduced peak angle within the C-start response. Further, the differences in kinematic changes across all three cave populations, raise the possibility that different genetic and neural mechanism underlie changes in this escape response across different population.

Identifying the behavioral and neuronal components of the C-Start response that are associated with effective interspecies and intraspecies escape may provide insight into the ecological factors contributing to the evolution of the C-start. In guppies, increased angular speed during the first phase of fast start escapes has been correlated with more effective predator evasion, raising the possibility that individual kinematic parameters contribute to successful predator avoidance (Walker et al, 2005). Interestingly, all cave population analyzed here exhibit decreased angular speed. Further, it is extremely likely that a quick latency increases the likelihood of successful evasion. While it may seem intuitive to predict that an increase in response probability would be beneficial in successfully avoiding predators, in a situation where predators are known to rely heavily on mechanosensory stimuli for prey capture, such as in cavefish, initiation of a startle response could potentially be detrimental (Lloyd et al, 2018; Yoshizawa et al, 2010). These data suggest that the C-start responses of cavefish may be less effective for successful predator evasion as a result of the relaxation of predation in the cave environment.

The escape response is likely to be energetically expensive, and therefore extremely detrimental in the nutrient-limited cave environment. A possible explanation for the increased response probability of Pachón larvae to vibrational stimuli may be related to a shift in feeding strategy. In hunting archer fish and goldfish, C-shaped flexions have been associated with prey capture (Wohl & Schuster, 2007; Canfield 2007). Furthermore, in goldfish, this feeding behavior has been correlated with firing of the Mauthner neurons (Canfield & Rose, 1993). In cave populations of *Astyanax*, loss of eyesight has resulted in a shift in prey capture behavior involving the use of the lateral line to sense prey, which are captured using a C-bend, similar to the C-start behavior we examine here. This is in contrast to sight-dependent prey capture observed in surface fish which consists of J-shaped turns and a of a head-on approach. Interestingly, Pachón cavefish were able to successfully capture prey even after complete pharmaceutical ablation of the lateral line, but were unable to capture dead prey, suggesting hat alternate modes of perception of movement are being utilized (Lloyd et al, 2018). Taken together, these data suggest that the increase in acoustically driven C-start responses observed in cavefish may be driven by a shift in feeding strategy.

Powerful genetic approaches in zebrafish have provided extensive mechanistic insight into the function of the Mauthner neurons (Burgess et al, 2014; Shimazaki et al, 2019; Stil & Drapeau, 2015; Monesson-Olson et al, 2014). This includes the use of genetically expressed Ca2+ sensors to identify how the activity of these neurons is modulated and the use of GAL4-based genetic screens to identify additional circuits that regulate the startle response (Takahashi et al, 2017; Lacoste et al, 2015; Choe et al, 2013). Recently, many of these technologies including GCaMP imaging, *tol2* transgenesis, and CRISPR gene editing have been developed in *A. mexicanus* (Stahl et al,2019; Kowalko et al, 2018; Elipot et al, 2014). The application of these genetic approaches has potential to define functional differences between surface fish and cavefish and provide mechanistic insight into evolved differences between the populations. For example, the anatomy and activity of Mauthner neurons can be directly compared between individual populations. Our identification of differences in response probability and kinematics between *A. mexicanus* populations position this system as a powerful model for examining the evolution of the escape responses and sensory-motor integration.

## Methods

Animal husbandry was carried out as previously described (Borowsky, 2008; Stahl et al, 2019) and all protocols were approved by the IACUC Florida Atlantic University (Protocols A15-32 and A16-04). Fish were housed in the Florida Atlantic University core facilities at 23 ± 1°C constant water temperature throughout rearing for behavior experiments (Borowsky, 2008). Lights were kept on a 14:10 hr light-dark cycle that remained constant throughout the animal’s lifetime. Light intensity was kept between 25–40. Larvae were raised in 200ml bowls.

### Behavioral experiments

C-start responses were elicited according to methods previously utilized for zebrafish (Burgess & Granato, 2007; Bhandiwad et al, 2013; Zeddies & Fay, 2005). All behavioral testing was done between ZT5 and ZT9 (Zeitgeber time) in a temperature controlled room maintained between 23 and 25°C. For all assays individual 6 dpf larvae were placed within 15×15×9 cm square wells on a custom 3D-printed polyactic acid plate (Autodesk Fusion 360; San Rafael, CA; Creality CR10 Max; Guangdong, China) and allowed to acclimate for 10 minutes before being exposed to a single stimulus. The plates were securely screwed onto a vertically oriented vibration exciter (Type 4810; Bruel and Kjaer, Duluth, GA) controlled by a multifunction I/O device (PCIe-6321; National Instruments, Austin, TX). Stimuli were 500 Hz square waves of 50 ms duration generated using Labview 2018 v.18.0f2 (National Instruments, Austin, TX) and were of an intensity of 31 dB, unless otherwise stated. Stimulus intensity was determined using a Check Mate CM-130 SPL meter (Galaxy Audio; Wichita, KS) held approximately 2 cm above the center of the vibrating apparatus. Plates had between 1 and 18 wells. For trials conducted on plates with greater than 6 wells, recording was done from above with the well placed directly over the center of the exciter to avoid shifting the center of mass away from the source of the stimulus. In these cases, lighting was provided from below using LED strips in addition to overhead ceiling lights. For trials conducted using plates with 6 wells or fewer, recording was done from below and illumination was done from above using LED strips and a polycarbonate sheet for diffusing light. Infrared light strips (940 nm) were used for all light/dark experiments. For trials conducted for the light condition, white light LED strips were also used.

Video was acquired at 1000 frames per second using an FPS 2000 high speed camera (The Slow Motion Camera Company Limited; London, UK).

### Analysis of C-start responses

C-start responses were identified as accelerated, simultaneous flexion of the head and tail in the same direction. Response probability is reported as the total proportion of larvae that exhibited a C-start response. Kinematic analysis was performed by separately analyzing various parameters of the C-start response as previously done in zebrafish (Issa et al, 2011; Burgess & Granato, 2007; Takahashi et al, 2017). The “angle” tool available on ImageJ 1.52a (National Institutes of Health; Bethesda, MD) was used to determine the orientation of the larvae by measuring the angle formed by a horizontal line and a line drawn along the midline of the fish from the anterior-most point of the swim bladder to the anterior-most point on the head. Measurements were subsequently standardized to the orientation of the larva 1 ms before stimulus onset.

Response latency was defined as the time between stimulus onset and a change in orientation of 10°. Peak angle was identified as the maximum change in orientation before a change in direction back toward the original orientation. Speed was determined as the slope of the best-fit line for change in body orientation from the point in time designated as the latency to the time of the peak angle.

### Statistical Analysis

All statistical tests were conducted on GraphPad Prism 8.2.1 or RStudio 1.2.1335. Differences in response probability were analyzed using Fisher’s Exact Test, except for analyses of 2×3 tables, which were analyzed using a X^2^ test. All error bars on response probability data denote margin of error of the sample proportion calculated using a z*-value of 1.96. Post-hoc analysis was conducted on results that indicated a significant difference (α ≤ 0.05) via pairwise X^2^ tests and Bonferroni correction of p-values. Normality of kinematic data was assessed using a Shapiro-Wilk test. Data that did not pass the normality test were subsequently assessed using the Mann-Whitney test and data that did pass the normality test were assessed using an unpaired t-test. In cases involving more than 2 populations, a one-way ANOVA was used followed by Tukey’s test in cases that the results of the ANOVA indicated significant differences (α ≤ 0.05). Correlation between kinematic parameters was assessed using Spearman’s rank order correlation.

